# Endoplasmic reticulum membrane contact sites coordinate exocytic site assembly and activity in neuroendocrine cells

**DOI:** 10.64898/2026.02.04.703764

**Authors:** L. Streit, F. Delavoie, A. Wolf, C. Royer, A.M. Haeberlé, S. Hugel, S. Gasman, N. Vitale, S. Chasserot-Golaz

**Affiliations:** Centre National de la Recherche Scientifique, Université de Strasbourg, Institut des Neurosciences Cellulaires et Intégratives, F-67000 Strasbourg, France; Molecular, Cellular and Developmental Biology Unit (MCD), Centre de Biologie Integrative (CBI), University of Toulouse, UPS, CNRS, Toulouse, France; Plateforme “Imagerie In Vitro” UAR3156 CNRS, Université de Strasbourg, France

**Keywords:** Calcium-regulated exocytosis, Endoplasmic reticulum, Membrane contact sites, Orai1, STIM1, Neuroendocrine secretion, Chromaffin cells

## Abstract

Membrane contact sites (MCS) between intracellular organelles regulate lipid exchange, organelles dynamics and spatial organization of signaling pathways, yet their contribution to regulated exocytosis remains poorly understood. Here, we investigated the role of endoplasmic reticulum (ER) MCS in calcium-regulated exocytosis using primary bovine chromaffin cells. Combining electron microscopy on plasma membrane (PM) sheets with immunogold labeling, we identified ER structures contacting docked secretory granules and classified three types of MCS: ER-PM, ER–granule and tripartite ER–PM–granule contacts. These contacts are enriched at exocytic sites and contain Orai1 and STIM1, both known for mediating store-operated calcium release. Functional perturbation of the Orai/STIM pathway revealed that constitutive STIM activation or pharmacological inhibition of Orai1 reduced the number of exocytotic events, slowed catecholamine release and disrupted actin organization at granule docking sites. Together, our findings revealed a previously unrecognized role for ER MCS in organizing exocytic sites and controlling secretion efficiency in neuroendocrine cells.

## Introduction

Calcium-regulated exocytosis is a fundamental cellular process in which secretory vesicles or granules fuse with the plasma membrane (PM) to release their contents, thereby supporting essential functions such as cell migration, wound repair, neurotransmission and hormone secretion ^1^. In neurons and neuroendocrine cells, calcium-dependent exocytosis has been extensively studied for decades, leading to the identification of many molecular components controlling secretory granule (SG) recruitment, docking and fusion with the PM ^2–4^. Among these, the SNARE complex has been identified as the minimal protein fusion machinery activated by calcium-sensors^5^. However, it is now clear that this protein machinery operates within a highly dynamic lipid environment, in which specific lipid species and their rapid remodeling actively contribute to exocytic site organization and function ^6, 7^. However, despite these advances, how exocytic sites are structurally organized to spatially coordinate vesicle tethering, assembly of the release machinery and fusion remains incompletely understood.

In neuroendocrine cells, SG tethering and docking to the PM occurs at specialized exocytic sites defined, in part, by distinct lipid microdomains ^8^. Electron tomography revealed that these domains underlie the bundling of actin filaments connecting SG to the PM, whereas functional experiments demonstrated the direct involvement of these actin bundles in granule recruitment and fusion ^9^. However, beyond actin bundles, tomographic analyses also suggested the presence of additional, yet uncharacterized, membrane structures associated with docked granules and the base of actin bundles close to the plasma membrane. These observations are consistent with earlier report describing endoplasmic reticulum (ER) structures in the immediate vicinity of SG at the PM of chromaffin cells, raising the possibility that cortical ER contributes to the establishment of functional exocytic sites^10^.

The ER forms membrane contact sites (MCS) with most cellular compartments, including lysosomes, endosomes, mitochondria, lipid droplets, and the PM. Through these contacts, the ER regulates lipid exchange, calcium fluxes and signaling events essential for cellular homeostasis ^11, 12^. ER MCS are highly dynamic structures that assemble and disassemble in response to physiological demands ^13^. Their formation relies on dedicated tether proteins that not only physically bridge membranes but also confer specific functional properties, including lipid transfer between membrane compartments. For instance, extended synaptotagmins (E-Syts) tether the ER to the PM in a calcium- and phospholipid-dependent, while also participating in phospholipid tranfer ^14^. Other tethers, such as Calcium Release-Activated Calcium Modulator 1 (Orai1) and STromal Interaction Molecule 1 (STIM1) regulate intracellular calcium homeostasis. STIM1 is an ER transmembrane protein sensitive to local calcium concentration, whereas Orai1 is a selective calcium channel located at the PM. Upon ER calcium depletion, STIM1 oligomerizes, accumulates at ER-PM contact sites and activates Orai1, enabling extracellular calcium entry and refilling of ER calcium stores ^15^. Importantly, in neurosecretory PC12 cells, Orai1 localizes to SG membranes, in contrast to its classical description as a PM channel ^16^.

Given the central role of calcium signaling and lipid organization at exocytic sites, together with the emerging function of ER MCS in intracellular signaling and lipid metabolism, we hypothesized that ER MCS may actively participate in the assembly and regulation of exocytic sites in neuroendocrine cells. Here, by using primary bovine chromaffin cells and combining transmission electron microscopy, electron tomography of PM sheets and functional perturbations of STIM1-Orai1 signaling, we identified and characterized dual and tripartite ER-PM-SG contact sites formed during stimulation and demonstrate their functional importance to regulated exocytosis.

## Results

### Cortical ER associates with plasma membrane and exocytic sites in chromaffin cells

Based on previous observations describing peripheral ER structures in close proximity SG ^10^, we asked whether the membrane structures detected near docked SG and at the base of actin bundles tethering SG to the PM ^9^ correspond to ER. We immunolabeled resting (unstimulated) and nicotine-stimulated chromaffin cells using anti-calnexin antibodies (ER marker) ^17^ and anti-SNAP25 antibodies (PM marker) ^18^. Following stimulation, calnexin exhibited a clear increased colocalization with SNAP25, indicating the recruitment of cortical ER to the PM (Fig. 1a). Semi-quantitative fluorescence analyses revealed that ∼45% of the calnexin signal associated with the PM in stimulated cells, compared with < 30% in unstimulated cells (Fig. 1b), consistent with the formation or reinforcement of ER-PM contacts during exocytosis.

**Fig. 1.**
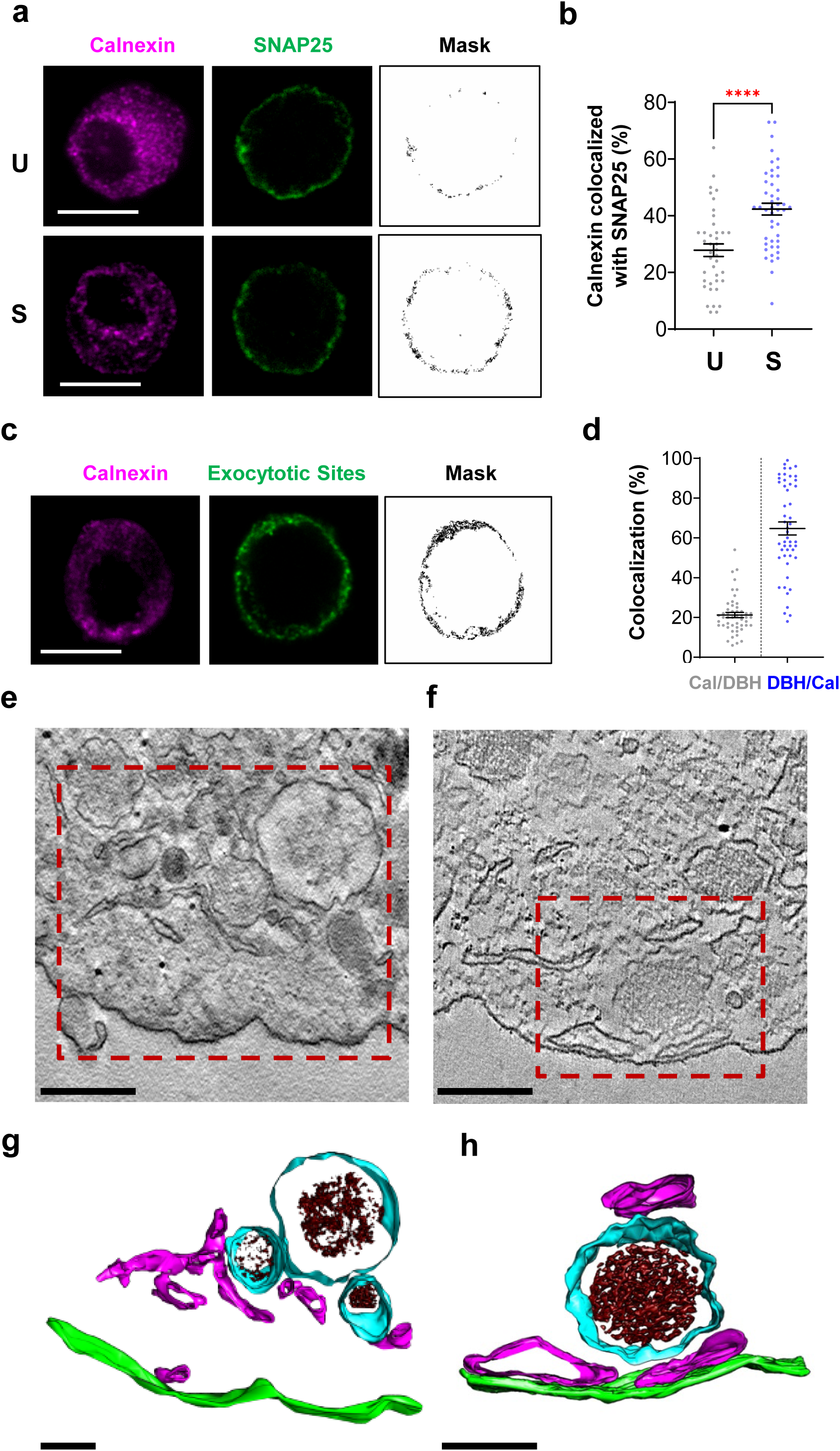
ER MCS form at exocytic sites upon chromaffin cell stimulation. **a** Confocal images of chromaffin cells left unstimulated (U) or stimulated with nicotine for 3 min (S), fixed, and immunolabeled for calnexin (magenta) and SNAP25 (green), scale bars: 10 µm. Masks were obtained by selecting dual labelled pixels with FIJI. **b** Quantification of calnexin colocalization with SNAP25 in unstimulated (gray) or stimulated cells (blue). Data are represented as mean ± SEM, each point represents a measure from a single cell; ****: *p* < 0.0001 (Mann-Whitney test; n ≥ 40 per condition from three independent experiments performed on different cultures). **c** Confocal images of labeled exocytic sites and ER, cells were stimulated with 20 µM nicotine for 3 min with anti-DBH antibodies, fixed, permeabilized, and stained for calnexin (magenta) and DBH (green), scale bar: 10 µm. **d** Quantification of colocalization between calnexin and exocytotic sites (DBH positive pixels, gray) or exocytic sites and calnexin positive pixels (blue). Data are represented as mean ± SEM, each point represents a measure from a single cell (n ≥ 50 per condition from two independent experiments performed on different cultures). **e**, **f** Examples of a tomographic section of unstimulated (e) or nicotine-stimulated (f) chromaffin cells, scale bars: 200 nm. **g, h** 3D surface rendering of the subtomogram of regions outlined in boxes in panels e and f (red dotted lines) showing ER (magenta), SG membrane (cyan), SG matrix (brown) and PM (green), scale bars: 100 nm.

To test whether this cortical ER specifically associates with exocytic sites, we immunolabeled ER (calnexin) in chromaffin cells stimulated in the presence of anti-Dopamine-β-Hydroxylase (DBH) antibodies, a luminal SG marker exposed only after fusion ^19^ (Figure 1c). Nicotine stimulation induced a peripheral punctate DBH pattern consistent with active exocytosis, and calnexin labeling frequently colocalized with DBH-positive spots (Figure 1c). Quantification of the masks revealed that approximately 20% of ER structures were positioned close to exocytic sites, whereas more than 60% of exocytic sites were contacted by the ER (Fig. 1d).

To visualize the ultrastructure of these potential contacts, we performed electron tomography on ultrathin sections from unstimulated (Fig. 1e, g, videos 1, 2) and nicotine-stimulated chromaffin cells (Fig. 1f, h, videos 3, 4). In resting cells, tomograms revealed ribosome-free ER domains positioned close to SGs, consistent with potential ER-SG contacts, whereas dual ER–PM associations were only occasionally observed. The space observed between SGs and the PM may reflect the presence of a filamentous actin barrier in resting chromaffin cells ^20^. To confirm that these ribosome-free structures corresponded to ER, we performed calnexin labeling on ultrathin sections of unstimulated and nicotine-stimulated chromaffin cells, validating their ER identity (Supplemental Fig. 1). Strikingly, upon stimulation, both ER and SG structures shifted toward the PM, dynamically establishing potential dual and even tripartite ER–SG–PM contacts. These ultrastructural observations are consistent with the increased calnexin/SNAP25 colocalization detected upon stimulation (Fig. 1a), supporting the concept that the ER forms dynamic MCS with the PM and SG during exocytosis.

### Ultrastructure of ER MCS at exocytic sites in chromaffin cell

Transmission Electron Microscopy (TEM) at high magnification of ultrathin sections of chromaffin cell showed close proximities between docked chromaffin granules and the ER (Fig. 2). Three different types of ER-dependent MCS were consistently observed i) ER-PM contacts (Fig. 2a), ii) ER-SG contacts (Fig. 2b), and iii) tripartite ER-SG-PM contacts (Fig. 2c).

**Fig. 2.**
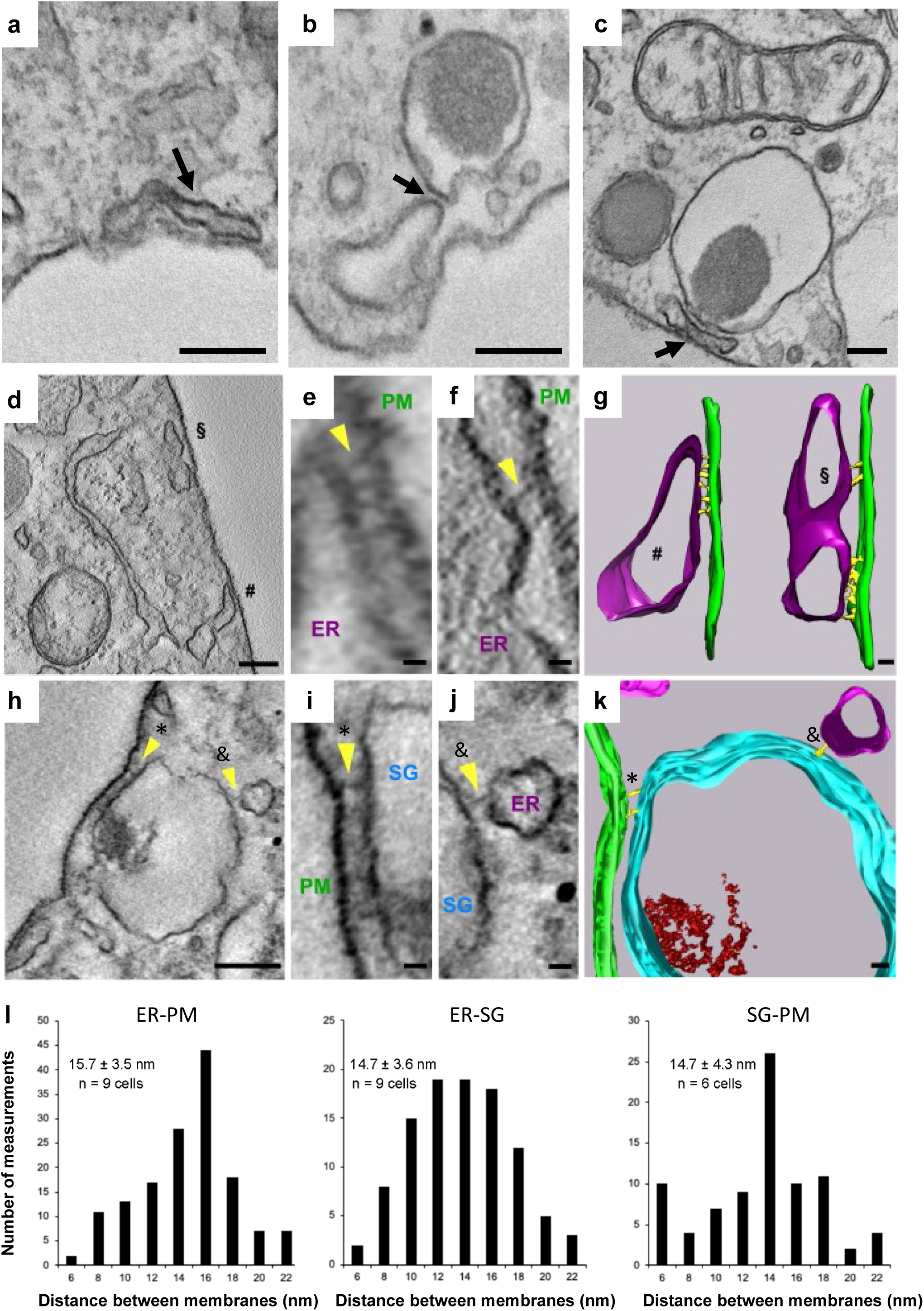
Ultrastructure of ER MCS at exocytic sites in stimulated chromaffin cells. a-c. Representative electron micrographs of chromaffin cells showing three distinct types of MCS upon 3 min nicotinic stimulation (black arrows): ER-PM (a), ER-SG (b) and tripartite ER-PM-SG (c), scale bar: 200 nm. **d** Tomographic section illustrating ER MCS with the PM, scale bar, 200 nm. **e** and **f** Enlarged views of the regions marked with # (e) and § (f) in panel d. Yellow arrows indicate fine filaments tethering both membranes compartments, scale bars: 10 nm. **g** 3D surface rendering of the tomogram of panels e and f showing ER (magenta), PM (green), and tethering filaments (yellow), scale bar: 20 nm. **h** Tomographic section illustrating ER MCS with SGs, scale bar: 200 nm. Yellow arrows indicate tethering filaments between SG and PM (*) and SG and ER (&). **i, j** Enlarged views of the regions marked with * (i) and & (j) in panel H, scale bars: 20 nm. **k** 3D surface rendering of the tomogram of the panel h showing ER (magenta), SG (cyan), PM (green), and tethering filaments (yellow), scale bar: 20 nm. **l** Distribution of intermembrane distances measured between ER-PM, ER-SG, and SG-PM contacts.

Next, we performed three-dimensional electron tomography combined with segmentation to improve visualization of ER MCS (Fig. 2d-k). The reconstructed 3D models confirmed the presence of the three types of MCS and revealed that SG outer membranes closely contacted both the ER and the PM without showing apparent signs of fusion (Fig. 2k). Importantly, we consistently detected short filamentous structure connecting the ER to either the PM or SGs (Fig. 2g, k), potentially corresponding to tether proteins. A morphometric analysis indicated that the length of these ER-PM and ER-SG tether filaments measured 15.7± 3.5 nm and 14.7 ± 3.5 nm, respectively (Fig. 2l). These distances match those reported for Ca^2+^-dependent E-Syt1-mediated ER-PM MCS ^14, 21, 22^ and for STIM1-dependent ER-PM MCS ^23, 24^. Together, these results support the involvement of specialized ER-mediated MCS with SGs and/or the PM in exocytosis. In addition, short filaments directly linking SGs to the PM were observed with an average length of 14.7 ± 4.3 nm (Fig. 2k, l), consistent with the dimensions of annexin A2 tetramers ^25^, which are key mediators of SG docking ^9^.

### Characterization and topography of ER MCS formed during exocytosis

To further characterize these ER MCS at the ultrastructural level, we performed immunogold-TEM on native PM sheets (Fig. 3). This method provides a two-dimensional view of the cytoplasmic leaflet of the PM along with its associated proteins. We have previously adapted this approach to chromaffin cells and neurons to visualize large pieces of PM containing embedded proteins and docked vesicles, enabling the identification of proteins present at exocytic sites by immunogold staining ^9, 26, 27^. Immunogold labelling of calnexin (10 nm gold particles) on PM sheets from unstimulated vs. nicotine-stimulated cells revealed striking differences. In unstimulated cells, calnexin was detected only in small moderately electron-dense structures (Fig. 3a, Unstimulated), consistent with the limited ER-PM contacts observed by EM (Fig. 1e, g, Supplemental Fig.1). Conversely, in stimulated cells calnexin positive structures were larger and concentrated near docked SG (Fig. 3a, Stimulated), in agreement with the tripartite contacts observed by tomography (Fig. 1h). Edge-to-edge distance measurements showed that nearly 75% of calnexin-positive ER structures were located within 50 nm of docked SG confirming a strong enrichment of ER MCS at exocytic sites (Fig. 3b).

**Fig. 3.**
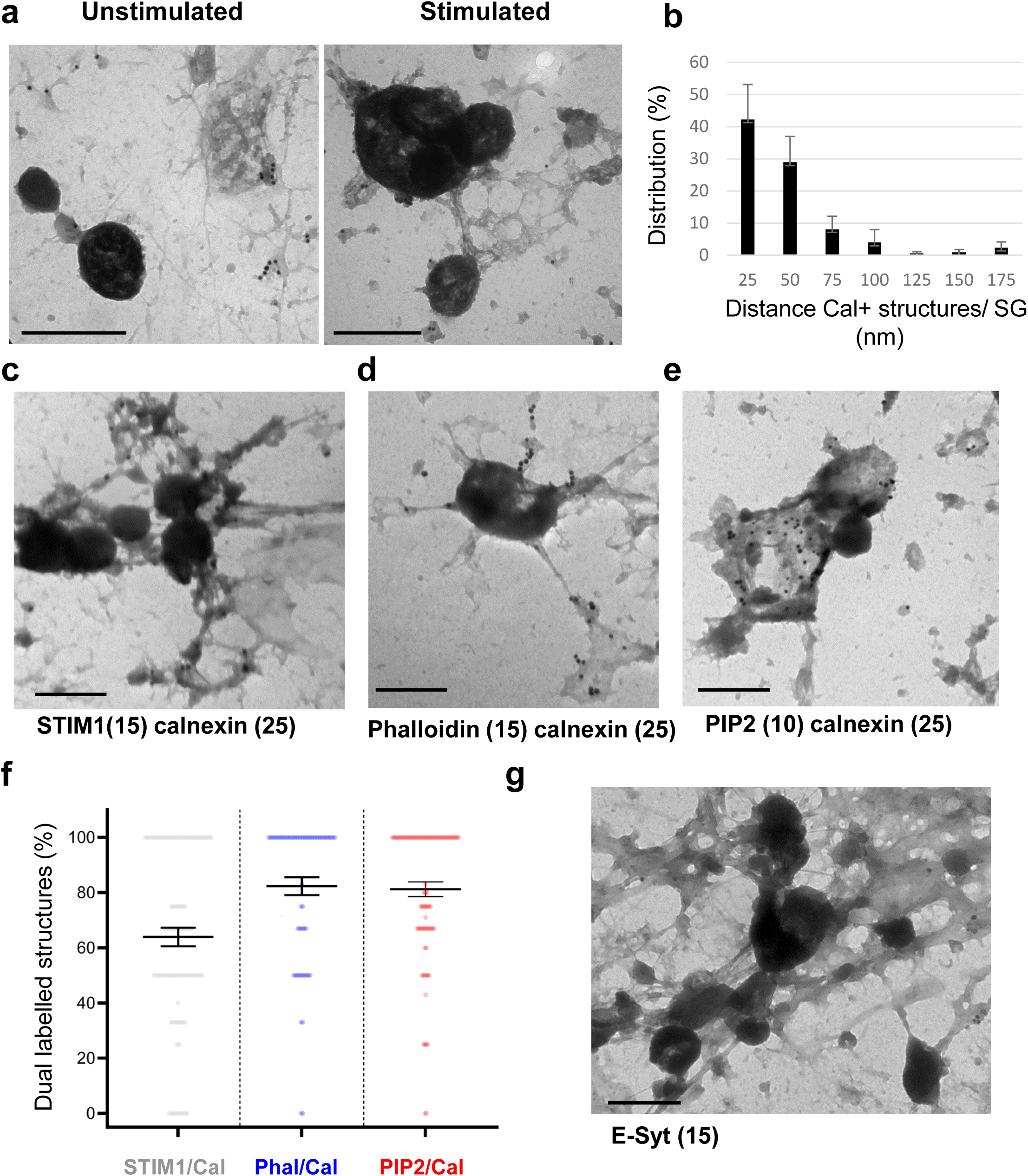
STIM1, actin filaments, PIP2 and E-Syt are present at MCS in stimulated cells. **a** TEM images of PM sheets from unstimulated or 3 min nicotine-stimulated chromaffin cells labeled for calnexin with 10 nm gold beads, scale bar: 500 nm. **b** Distribution of calnexin-positive structures relative to the edge-to-edge distance of docked granules. Data are represented as mean ± SEM (n = 120 measures from 5 independent experiments performed on different cultures). Distances were measured manually with FIJI. Calnexin-positive structures were enriched within 25 nm of granule edges. **c - e** Representative TEM images of stimulated PM sheets labeled for calnexin (25 nm gold beads) and STIM1 (15 nm gold beads, c), F-actin (Phalloidin, 15 nm gold beads, d) or PIP2 (10 nm gold beads, e), scale bar: 500nm. **f** Percentage of calnexin-positive membrane patches (ER MCS) associated with STIM1 (gray), phalloidin (blue) or PIP2 (red) beads. Quantifications were performed manually in Photoshop. Data are represented as mean ± SEM with each point representing a measure from a single image (n = 91; 65 and 89 respectively per dual staining from three independent experiments performed on different cultures). **g** Representative TEM image of stimulated PM sheets labeled for E-Syt (15 nm gold beads), scale bar: 500 nm.

Next, to better characterize the molecular organization of ER MCS, we performed immunogold labeling of calnexin together with STIM1, actin filaments or PIP2. The spatial distribution of distinct gold particle sizes revealed a clear co-localization of STIM1 and calnexin within the same structures, indicating that STIM1 is a prominent component of ER MCS. Quantitative analyses showed that approximately ∼70% of calnexin-positive structures were also STIM1-positive (Fig. 3c, f), consistent with the idea that ER MCS correspond to specialized ER subdomains positioned at ER-PM interfaces.

Based on our previous observations of actin bundles at exocytic sites ^9^ and on earlier reports indicating that cortical actin contributes to the spatial organization of ER–PM junctions ^28^, we next examined the association between ER MCS and the actin cytoskeleton. Dual labeling of calnexin and filamentous actin using biotinylated phalloidin (15-nm gold) revealed dense filamentous actin structures connecting docked SGs to the PM, as well as actin enrichment at the periphery of calnexin-positive ER domains (Fig. 3d). Quantitative analysis showed that ∼80% of ER MCS were associated with F-actin in proximity to docked SGs (Fig. 3f), suggesting that actin filaments may contribute to the spatial organization and stabilization of ER-PM contact sites at exocytic regions.

Because ER-PM MCS are established sites for lipid exchange between membrane compartments ^13^, we next examined the distribution of phosphatidylinositol 4,5-bisphosphate (PIP2), a key lipid for exocytosis ^29, 30^. PIP2 labeling was detected in approximately ∼ 80% of ER MCS (Fig. 3e, f), with a marked enrichment at ER-PM MCS (∼79%,). Consistently, immunogold labeling of plasma membrane sheets revealed that E-Syt, a known ER–PM lipid transfer protein, was predominantly localized at ER-PM MCS (Fig. 3g, ∼87%), in agreement with a role for these sites in local PIP2 exchange between the ER and the PM.

Altogether, these data indicate that ER MCS in secretory cells represent highly organized membrane domains enriched in STIM1 and E-Syt, aligned with PIP2-rich plasma membrane regions and closely associated with an F-actin scaffold. This organization supports a model in which ER MCS participate to local calcium regulation, lipid exchange and contribute to the spatial stabilization of secretory granule docking at defined exocytic sites along the plasma membrane.

We further used electron tomography of PM sheets to resolve the spatial architecture of ER structures formed at the cytoplasmic face of the PM in nicotine-stimulated cells (Fig. 4). As suggested by immunogold analyses (Fig. 3a), calnexin-positive ER structure formed direct contacts with the PM and, in doing so, also connected SG to the PM (Fig. 4a, b, g, h, videos 5-8). Three-dimensional reconstructions (Fig. 4c-f, i-k) further revealed that the ER adopts a distinct topography around docked SG, confirming that ER MCS are specifically integrated into exocytic sites architecture. Notably, actin bundles linking SGs to the ER (black arrows) and to the PM were also observed (Fig. 4i). Taken together, these results indicate that secretagogue-induced stimulation triggers the formation of calnexin-positive ER structures at the PM, where they associate with both actin filaments and docked SGs. These findings support the idea that the ER and filamentous actin cooperate to organize the exocytic machinery, thereby shaping exocytosis in neuroendocrine cells.

**Fig. 4.**
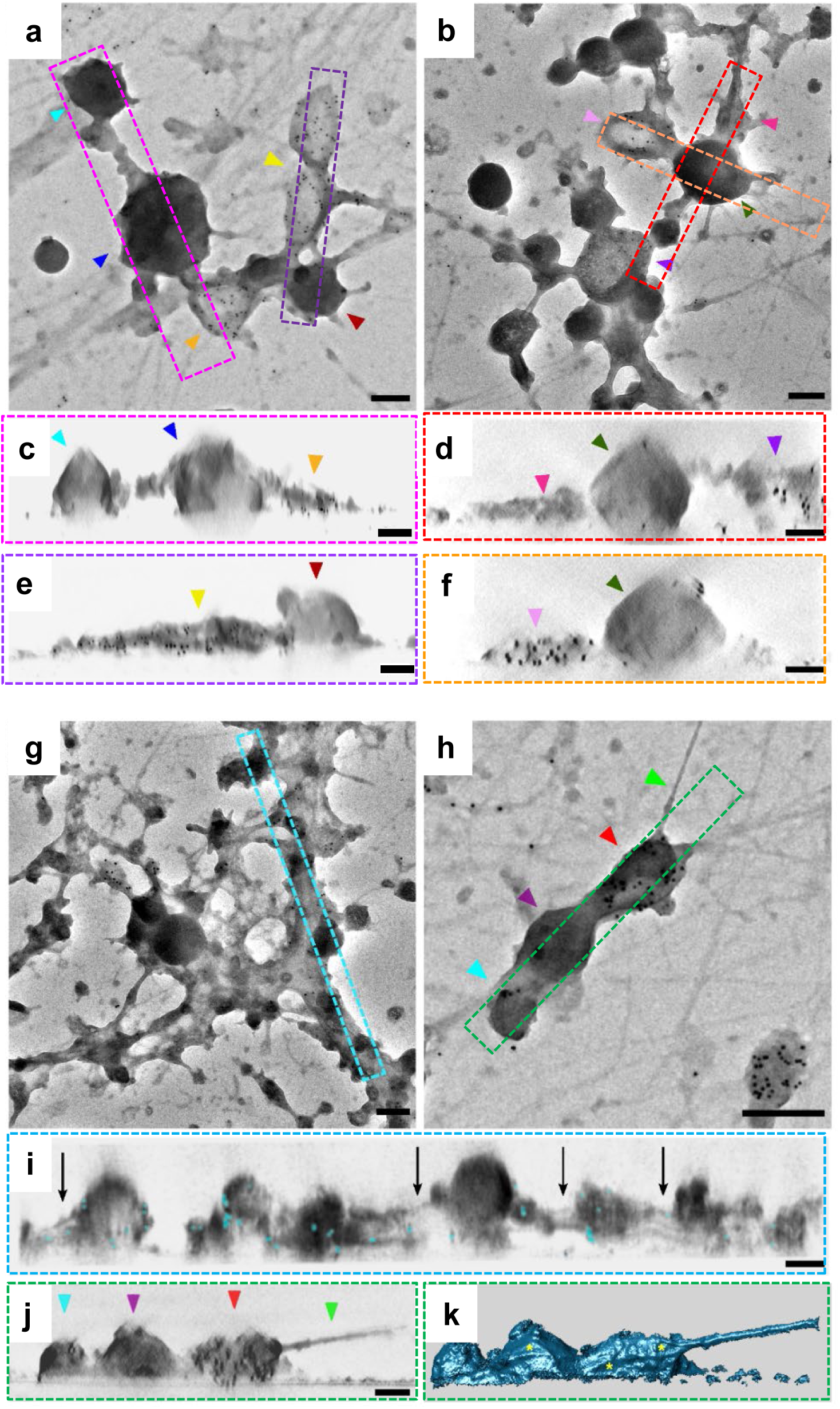
Topological organization of ER MCSs at exocytic sites. **a**, **b**, **g**, **h** Zero-tilt TEM images of calnexin-positive structures associated with SGs docked at the PM, scale bars: 200 nm. **c**-**f**, **i, j** Side views of isosurface representations from subtomograms corresponding to boxed regions in a, b, g, and h, scale bars: 100 nm. Colored arrowheads indicate landmarks for comparison between zero-tilt images and subtomograms. Black arrows highlight actin filaments in i. **k** Surface-rendered view of the subtomogram shown in j, yellow asterisks highlight actin filaments.

### Presence of STIM1 and Orai1 at MCS between ER and chromaffin granules

We next investigated whether ER MCS formed during exocytosis contain STIM1 and Orai1, two well-established markers of ER-PM MCS and central components of the calcium entry machinery ^31^. STIM1 is found exclusively at the ER membrane, whereas Orai1 has been described to both the PM and SG in PC12 cells ^16^, suggesting a potential role in coupling local calcium influx to exocytic events.

First, to determine whether STIM1 localizes at exocytic sites, we examine its distribution relative to DBH-positive fusion spots, hallmarks of transient post-fusion sites at the plasma membrane. STIM1 labeling showed clear colocalization with DBH, as confirmed by the mask image (Fig. 5a). Semi-quantitative analysis revealed that approximately 20% of STIM1 overlapped with exocytic sites (Fig. 5b), a value comparable to that observed for calnexin (Fig. 1c, d), suggesting that STIM1 is recruited to ER MCS formed at exocytic sites.

**Fig. 5.**
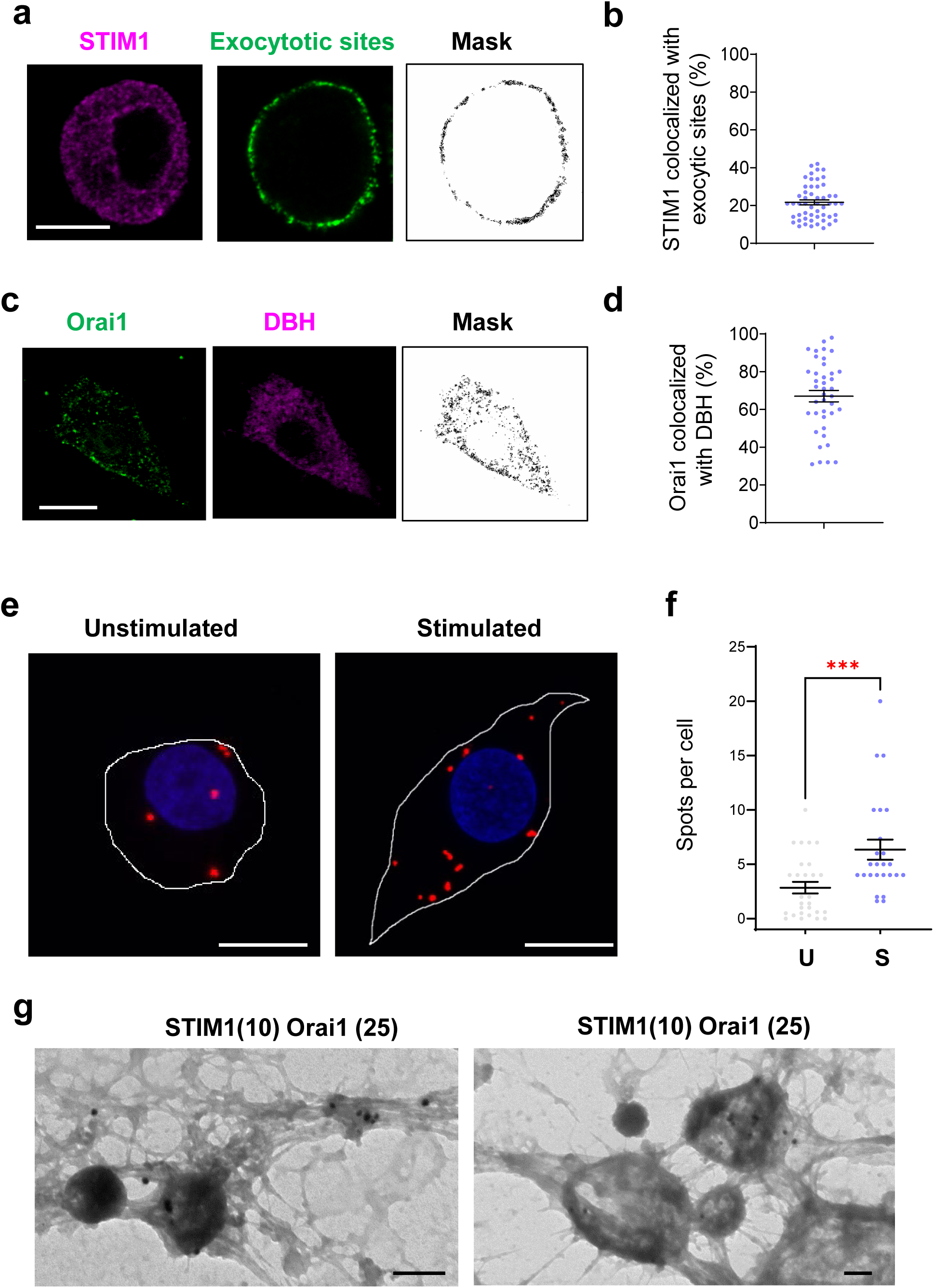
Orai1 is localized at the surface of chromaffin granules and interacts with STIM1 upon stimulation. **a** Confocal images of nicotine stimulated chromaffin cells (3 min) with anti-DBH antibodies, stained for STIM1 (magenta) and DBH (green). Masks were obtained by selecting dual labelled pixels with FIJI, scale bar: 10 µm. **b** Percentage of STIM1 colocalizing with DBH-positive exocytotic sites. Data are represented as mean ± SEM, each point represents a measure from an individual cell (n = 52 cells from three independent experiments performed on different cultures). **c** Confocal images of chromaffin cells stained for Orai1 (green) and DBH (magenta). Masks were obtained by selecting dual labelled pixels with FIJI, scale bar: 10 µm. **d** Percentage of Orai1 colocalized with DBH-positive SG. Data are represented as mean ± SEM, each point represents a measure from an individual cell (n = 40 cells from three independent experiments performed on different cultures). **e** Z projection of *In situ* proximity ligation assay (PLA) to probe STIM1 and Orai1 interaction in unstimulated (U) and stimulated (S, 3 min with nicotine) chromaffin cells. PLA signals appear as fluorescent puncta; nuclei (blue) were stained with Hoechst, scale bars: 10 µm. **f** Number of PLA spots detected per cell unstimulated (gray) and stimulated (blue). Data are represented as mean ± SEM, each point represents a measure from an individual cell; ***: *p* < 0.001 (Mann-Whitney test; n ≥ 25 images from two independent experiments performed on different cultures). **g** TEM image of PM sheets of stimulated cells dual-labeled for STIM1 (10 nm gold beads) and Orai1 (25 nm gold beads), scale bars: 200 nm.

Next, we assessed the presence of Orai1 on chromaffin SGs. Immunolabeling with anti-Orai1 and anti-DBH antibodies revealed a clear colocalization, indicating that Orai1 is present on SG (Fig. 5c). Quantification showed that ∼70% of Orai1 signal was associated with the SGs (Fig. 5d). To examine whether Orai1 and STIM1 come into close proximity during exocytosis, we performed an *in situ* Proximity Ligation Assay (PLA) ^32^ using antibodies against Orai1 and STIM1 (Fig. 5e, f, videos 9, 10). PLA generate fluorescent puncta only when two epitopes lie within ∼40 nm. Resting cells displayed an average of 3 fluorescent puncta per cell, indicating poor STIM1-Orai1 interaction. However, nicotine stimulation more than doubled the average number of fluorescent spots suggesting a significant increase in STIM1-Orai1 interaction during exocytosis and supporting the formation of functional ER-MCS with SG at exocytic sites.

Finally, dual immunogold-labelling of Orai1 and STIM1 on PM sheets from stimulated chromaffin cells also demonstrated their coexistence both at ER-SG and ER-PM MCS (Fig. 5g). Counting of beads revealed that 80 ± 2% of the STIM1-labeled beads were present on Orai1-positive structures, and 49 ± 1% were found on SG (85 images from 2 independent experiments). These observations suggest that STIM1-Orai1 complexes are formed at exocytic sites, where they may contribute to local calcium signaling required for granule exocytosis. To probe the functional role of this interaction, we next employed two complementary molecular and pharmacological strategies.

### Expression of the constitutively active STIM1-L251S mutant inhibits exocytosis

To probe the functional relevance of the STIM1/Orai1 complex, we overexpressed GFP as a reporter and either STIM1-WT or the constitutively active STIM1-L251S mutant in primary chromaffin cells. The L251S mutation locks STIM1 in an activated conformation that interacts more efficiently with Orai1 than the WT protein ^33–35^. Catecholamine release was then investigated using carbon fiber amperometry, which resolves both the number and kinetics of individual exocytic events ^36^. Fig. 6a shows representative amperometric traces recorded from chromaffin cells expressing GFP as a control, STIM1-WT or STIM1-L251S and Table 1 summarizes amperometric parameters. The constitutively active STIM1-L251S mutant markedly reduced the number of fusion events whereas the overexpression of STIM1-WT had no significant effect (Fig. 6b). Importantly, beyond modulating number of events, STIM1-L251S expression slowed catecholamine release kinetics (Fig. 6c, d). This was reflected by a decrease in spike amplitude, used as a proxy for maximal catecholamine flux. At the same time, we observed that STIM1-L251S expression decreased spike half-width (T_1/2_) and time-to-peak (T_peak_), parameters reporting on fusion-pore dilation and/or granule matrix expansion^37^. It is also noteworthy that overexpression of STIM1-WT was associated with similar changes in these kinetic spike parameters, albeit to a lesser extent; this observation could potentially be related to the reported effects of STIM1 overexpression on Orai1 distribution ^38, 39^. Together, these functional experiments demonstrate that the timing of STIM1 activation is critical for determining the number of vesicles that undergo fusion suggesting that STIM1 activation interferes with an early trafficking step upstream of fusion such as SG biogenesis, maturation transport and/or docking ^16^. Conversely, alteration in release kinetics seen with WT and constitutively active form of STIM1 appear to depend primarily on STIM1 abundance rather than on its activation state, suggesting a mechanistic link between STIM1/Orai1 channels activity through the establishment of MCS and the control of catecholamine flux through the fusion pore.

**Fig. 6.**
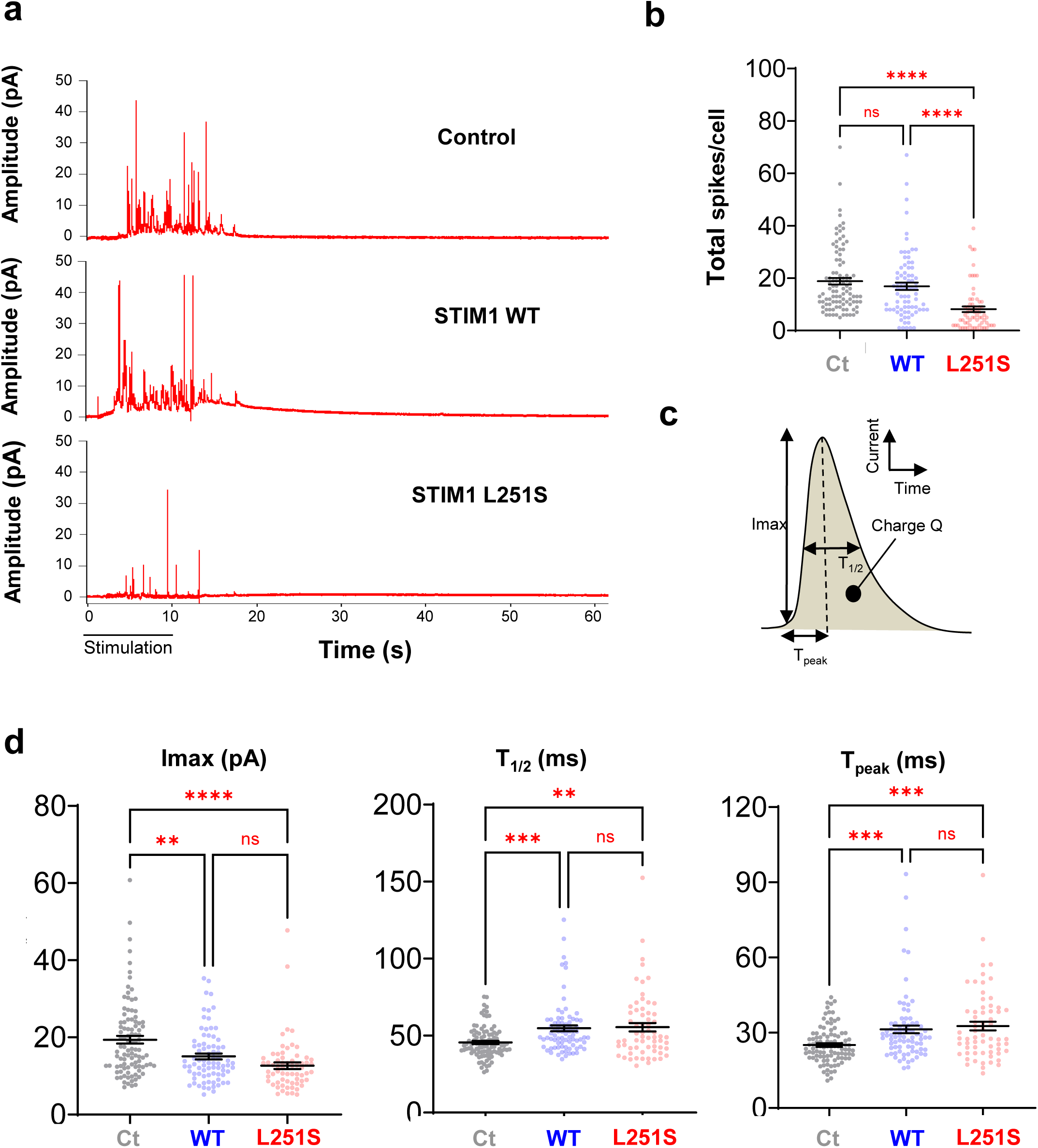
STIM1 mutant L251S overexpression impairs the number of fusion events and slows down their kinetics. Individual bovine chromaffin cells were stimulated by a local 10 s puff of 100 mM K⁺, and catecholamine secretion was monitored using carbon fiber amperometry. **a** Representative amperometric recordings of GFP (Control), STIM1-WT-GFP, or STIM1-L251S-GFP transfected cells. **b** Amount of amperometric spikes produced per cell. Data are represented as mean ± SEM, each point represents a measure from an individual cell (Gray: control, blue WT and red: L251S mutant); ns: non-significant and ****: *p* < 0.0001 (Kruskal-Wallis followed by Dunn’s multiple comparisons test; n ≥ 67 cells per conditions from three independent experiments performed on different cultures). **c** Schematic representation of individual spike parameters analyzed: quantal size (Charge Q), maximal current (I_max_), half-width (T_1/2_), and rise time (T_peak_). **d** Quantification of respectively, individual spike maximal current, half-width and rise time. Data are represented as mean ± SEM, each point represents the mean measure from an individual cell. ns: non-significant **: *p* < 0.01, ***: *p* < 0.001 and ****: *p* < 0.0001 (Kruskal-Wallis followed by Dunn’s multiple comparisons test; n ≥ 67 cells per conditions from three independent experiments performed on different cultures). Additional quantified parameters are shown in Table I.

**Table I:** Characteristics of amperometric spikes from bovine chromaffin cells transfected with GFP, STIM-WT or STIM-L251S mutant. Amperometric recordings were performed on cultured bovine chromaffin cells. Cells were transfected with GFP (control), STIM-WT, or STIM-L251S mutant. Cells were stimulated with a 10 s puff of 100 mM of K+. The number of recorded cells, amperometric spikes per cell, total analyzed spikes and single-event kinetic parameters are indicated. Data are expressed as mean ± SEM. **: p< 0.01, ***: p< 0.001 and ****p<0.0001 compared to control (Kruskal-Wallis followed by Dunn’s multiple comparisons test, data from three independent experiments performed on different cultures).

| Conditions | Control | STIM-WT | STIM-L251S |
| --- | --- | --- | --- |
| Number of cells | 98 | 79 | 67 |
| Total number of spikes | 1845 | 1335 | 546 |
| Number of spikes per cell | 18.83 ± 1.24 | 16.90 ± 1.41 | 8.15 ± 1.05 **** |
| Number of analyzed spikes | 1648 | 1194 | 466 |
| Charge Q (pC) | 0.90 ± 0.04 | 0.86 ± 0.05 | 0.72 ± 0.06 ** |
| I<sub>max</sub> (pA) | 19.39 ± 0.98 | 15.06 ± 0.76 ** | 12.67 ± 0.86 **** |
| T<sub>1/2</sub> (ms) | 45.58 ± 1.02 | 54.73 ± 1.93 *** | 55.47 ± 2.70 ** |
| T<sub>peak</sub> (ms) | 25.08 ± 0.72 | 31.37 ± 1.53 *** | 32.68 ± 1.75 *** |

### Inhibition of Orai1 with BTP2 impairs exocytosis and alters actin organization at docked SG

As a complementary approach, we treated the cells with 3,5-bistrifluoromethyl pyrazole (BTP2), a selective inhibitor of Orai1 ^40^ ^41^. As anticipated if Orai1 activity is important for regulated exocytosis, BTP2 treatment strongly reduced catecholamine secretion induced by either 10 µM nicotine or 59 mM K^+^ (Supplemental Fig. 2a). Because BTP2 can interfere with store-operated calcium channels ^40^, we next examined its effects on secretagogue-induced intracellular calcium signal in chromaffin cells. Interestingly, we observed that BTP2 reduced the calcium wave induced by nicotine (Supplemental Fig. 2b), whereas it did not affect calcium increase triggered by K^+^ depolarization (Supplemental Fig. 2c). Thus BTP2 treatment had no effect on voltage-gated channels activated by elevated extracellular K^+^ concentrations, as reported previously ^42^, suggesting that the inhibition of catecholamine secretion induced by BTP2 under K^+^ depolarization is independent of an effect on voltage-gated channels.

Carbon fiber amperometry revealed that BTP2 treatment decreased drastically the number of the amperometric events (Fig. 7a, b) and slowed catecholamine release kinetics (Fig. 7c). This was reflected by decreased spike amplitude together with increased half-width and time to peak (Table II). Notably, these effects closely mirrored those observed upon expression of the constitutively active mutant STIM1-L251S, and to a lesser extent of STIM1-WT (Fig. 6). This led us to conclude that impairing (either constitutive activation or inhibition) STIM1/Orai1 channels slowed down exocytosis, confirming the functional importance of those tethers in regulated exocytosis especially during SG fusion.

**Fig. 7.**
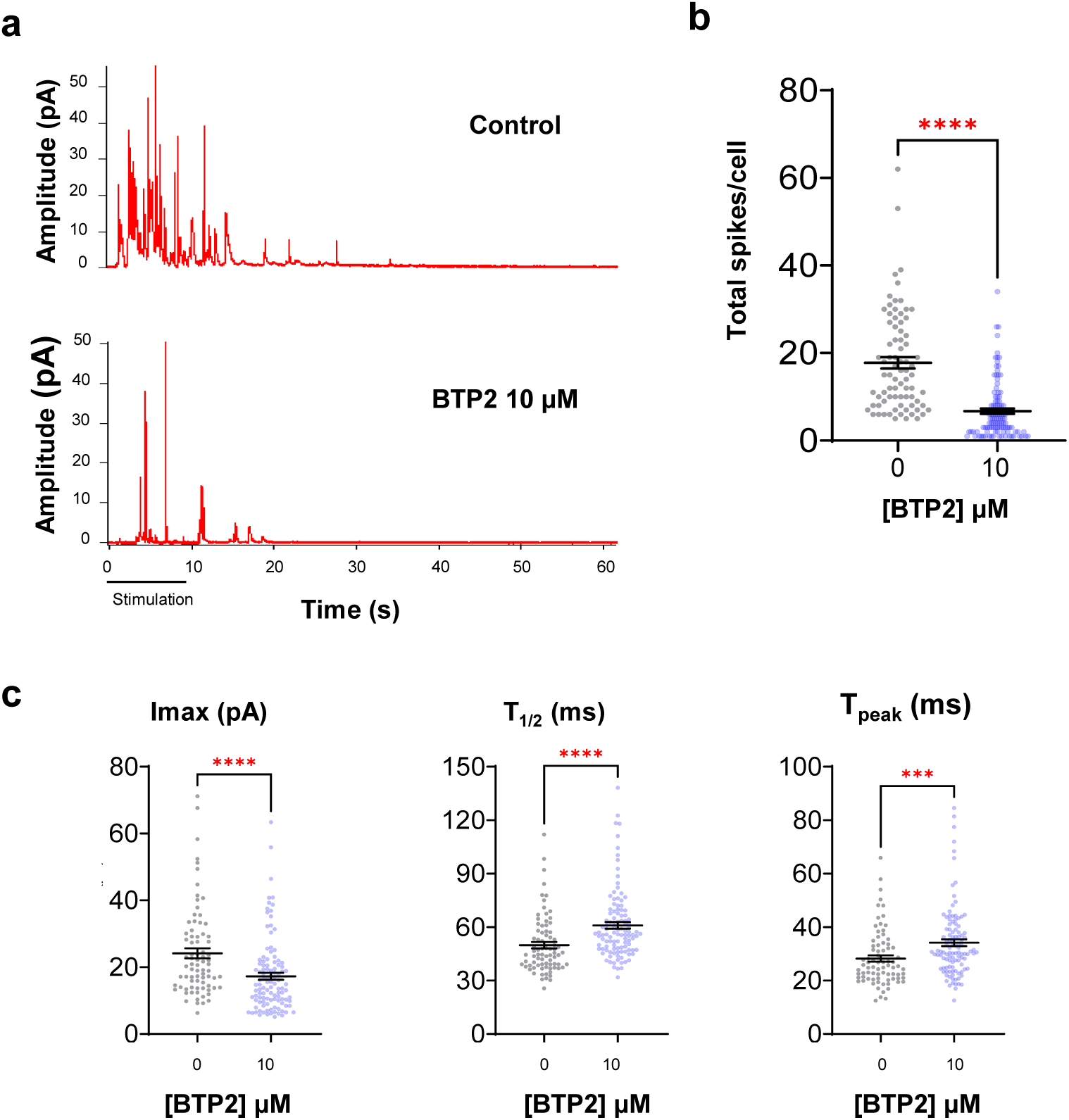
BTP2 decreases the number of fusion events and slows down fusion kinetics in chromaffin cells. Bovine chromaffin cells were incubated for 10 min with DMSO (control) or BTP2 (10 µM) and stimulated with a 10 s puff of 100 mM K⁺ to monitor catecholamine secretion using carbon fiber amperometry. DMSO and BTP2 remained in the bath during recording. **a** Representative amperometric recordings from control and BTP2-treated cells. **b** Quantification on the number of fusion events per cell. **c** Analysis of individual spike parameters (respectively maximal current, half-life and rise time) of treated cells. Data are represented as mean ± SEM, each point represents the mean measure from an individual cell (Gray: control and blue: BTP2). ***: *p* < 0.001 and ****: *p* < 0.0001 (Mann-Whitney test; n ≥ 79 cells per condition from three independent experiments performed on different cultures). Additional quantified parameters are shown in Table II.

**Table II:** Characteristics of amperometric spikes from bovine chromaffin cells treated with BTP2. Amperometric recordings were performed on cultured bovine chromaffin cells. Cells were pre-incubated 10 min with BTP2 10 µM and the molecule was present in bath during the amperometric recording. Cells were stimulated with a 10 s puff of 100 mM of potassium. The number of recorded cells, amperometric spikes per cell, total analyzed spikes, and single-event kinetic parameters are indicated. Data are expressed as mean ± SEM. ***: p<0.001 and ****: p< 0.0001 compared to control (Mann Whitney Rank Sum test, data from three independent experiments performed on different cultures).

| **[BTP2] μM** | **0** | **10** |
| --- | --- | --- |
| Number of cells | 79 | 110 |
| Total number of spikes | 1403 | 739 |
| Number of spikes per cell | $17.76 \pm 1.28$ | $6.72 \pm 0.62$ **** |
| Number of analyzed spikes | 1185 | 610 |
| Charge Q (pC) | $1.19 \pm 0.07$ | $1.06 \pm 0.06$ |
| I<sub>max</sub> (pA) | $24.14 \pm 1.50$ | $17.27 \pm 1.04$ **** |
| T<sub>1/2</sub> (ms) | $49.82 \pm 1.82$ | $60.95 \pm 1.84$ **** |
| T<sub>peak</sub> (ms) | $28.21 \pm 1.18$ | $34.15 \pm 1.26$ *** |

To identify which step of exocytosis is impaired by Orai1 inhibition, we performed PM sheets of chromaffin cells stimulated with high K^+^. BTP2 treatment did not visibly alter the number of docked SG, indicating that Orai1 activity is dispensable for granule docking (Fig. 8a, b). However, BTP2 treatment markedly reduced the density of actin filaments associated with SG, which appeared smoother than in untreated cells (Fig. 8b). In addition, the number of ER-derived reticular membranes (identified here as ER MCS) associated with docked SG was significantly reduced (Fig. 3), suggesting that Orai1 activation might also play structural roles in establishing MCS between the ER, the PM and/or SG.

**Fig. 8.**
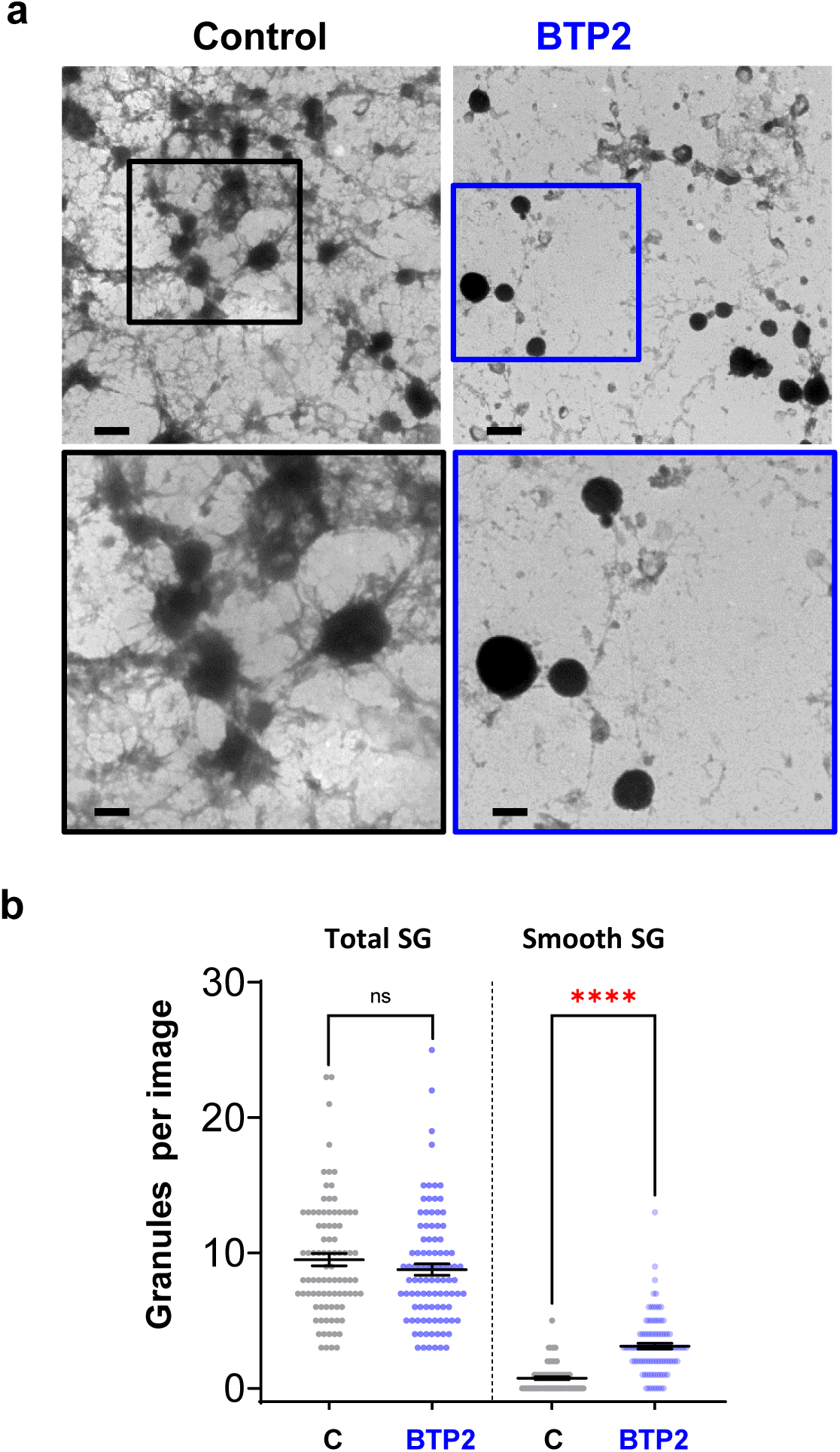
BTP2 impairs the formation of actin bundles linking secretory granules to the PM without altering the number of docked granules. **a** TEM image of PM sheets from stimulated chromaffin cells (3 min with 59 mM K^+^) that were untreated (control) or treated with BTP2 (10 µM), scale bars: 500 nm. Lower panels show enlarged views outlined in boxes in upper panels, scale bars: 250 nm. **b** Number of morphologically docked SG at the PM and smooth docked SG per image in control and BTP2-treated cells. Data are represented as mean ± SEM, each point represents a measure from an image. ns: non-significant and ****: *p* < 0.0001 (Mann-Whitney test; n ≥ 79 cells per condition from five independent experiments performed on different cultures).

Altogether, these results strongly support the idea that STIM1-Orai1-dependent MCS regulate both the organization of exocytotic sites and the efficiency of catecholamine release. In other words, our data suggests that STIM1-Orai1 found at the ER-PM MCS acts a modulator of exocytosis by stabilizing exocytotic sites by regulating local actin dynamics and Ca²⁺ concentrations in close proximity to chromaffin granules undergoing fusion while at the same time decreasing intraluminal SG calcium concentrations.

## Discussion

The objective of this study was to determine whether ER MCS are established at exocytosis sites and to assess their potential functional relevance for neurosecretion in primary chromaffin cells. The ER is widely recognized as the main cellular compartment responsible for lipid and protein biosynthesis and as the largest intracellular calcium reservoir. Owing to its extensive distribution throughout the cytoplasm, the ER forms MCS with different organelles and the PM, thereby regulating multiple cellular mechanisms ^11^. Our previous work revealed the three-dimensional organization of exocytotic site and identified previously uncharacterized membrane structures associated with SGs anchored to the PM via large actin bundles modulated by Annexin A2 ^9^. Given previous observation of ER structure near SG at the cell periphery in neuroendocrine cells ^10^, we tested whether these undefined membrane structures could be ER establishing MCS involved in the control of neurosecretion.

### Structural insights into cortical ER organization

Our study demonstrates that the ER establishes specialized MCS at exocytic sites in chromaffin cells. Indeed, in addition to the classical ER-PM contacts, ER-SG and tripartite ER-PM-SG contacts were also identified, especially in response to cell stimulation (Figures 1-3). A major strength of our approach is the use of endogenous markers for the ER (calnexin) and PM (SNAP25) combined with high-resolution ultrastructural and three-dimensional analyses. In contrast to other studies relying on overexpression of fluorescently tagged tethers, which can profoundly influence the size and dynamics of MCS ^14, 43^. Hence our strategy allowed us to visualize native contact sites with as little as possible perturbation of their organization. These experiments provide structural evidence that docked SG can physically engage with the cortical ER, thereby suggesting that the scope of these membrane contact sites may extend beyond classical ER–PM junctions. Furthermore, morphometric analysis revealed that the tethers between the different compartments displayed an average size of 15 nm (Fig. 2l), compatible with known MCS tethers. Additionally, our results are also in line with the recent observation that ribosome-free ER is in contact with insulin containing SG in β-cells ^44^, another type of neurosecretory cells.

We further demonstrate that ER–PM contacts increase upon stimulation, consistent with the dynamic nature of membrane contact sites observed across diverse cellular contexts^45^. These findings support the idea that the cortical ER forms a dynamic meshwork that contributes to secretory function. The E-Syt and PIP2 labellings on PM sheets suggest functional diversification among ER MCS, and implicates ER–PM contacts in the spatial organization of PIP2 at exocytic sites. By shaping local calcium and lipid microenvironments, ER–PM MCS may influence the fusion competence of docked granules. More broadly, our observations reinforce the emerging view that MCS act as dynamic signaling hubs, rather than passive structural interfaces ^46, 47^.

### Functional role of Orai1 on SG

A key functional advance of our study is the identification of the STIM1/Orai1 channel at ER-granule contact sites. Importantly, Orai1 localizes to chromaffin SG membranes, in contrast to its classical description as a PM channel ^16^. Its increased interaction with ER-resident STIM1 upon stimulation (Fig. 4) suggests a possible contribution in regulated exocytosis. Interestingly, in line with this hypothesis, previous studies reported stimulation-induced efflux of granular calcium in chromaffin cells ^48^. Accordingly, pharmacological inhibition of Orai1 with BTP2 strongly slowed the kinetics of individual fusion events upon cell depolarization (Fig. 7), despite leaving voltage-gated calcium entry intact (Fig. S2). At a mechanistic level, this could be explained either by alterations of fusion pore diameter triggered by less SNARE pin zippering due to a lower concentration of cytoplasmic calcium or changes in the diffusion of catecholamines from the granular matrix, due to higher luminal calcium concentrations^49, 50^. This latter hypothesis raises the possibility of a downstream mechanism in which impaired calcium release from granules may be associated with limited matrix expansion and reduced catecholamine dissociation from chromogranin A⁴⁹. This, in turn could be linked to inefficient ER refilling via retrograde coupling with the sarco/endoplasmic reticulum Ca²⁺-ATPase (SERCA) ⁵⁰ ⁵¹.

Consistent with earlier work that linked granular calcium to matrix compaction and maturation ^37, 51, 52^, these findings support a model in which STIM1/Orai1-mediated ER-granule contacts orchestrate calcium exchange directly at the organelle interface. This coupling ensures that SG are, not only properly matured, but also positioned within calcium-rich microdomains favoring exocytosis. Moreover, the loss of actin structures near SG upon BTP2 treatment (Fig. 7) suggests that local calcium release may promote polymerization of actin filaments at docking sites, thereby providing a structural scaffold required for stable granule attachment to the PM ^20, 53–55^.

### Integration of calcium and lipid signaling

Beyond calcium, ER-PM contacts are also well-known sites of lipid transfer between membrane compartments. Accordingly, lipid synthesis has been shown to control organelle biogenesis and dynamics ^56^. For instance, lipid exchange at these junctions is essential to maintain pools of PIP2 at the PM, which recruit priming factors such as syntaxin-1, organizing docking/priming in PIP2 enriched domains ^30, 57, 58^. These contacts may also supply phosphatidylserine (PS), which promotes calcium-dependent binding of synaptotagmin, lowering the energy barrier for membrane fusion ^59, 60^. In addition, phosphatidic acid (PA), another signaling lipid, playing pivotal roles in neurosecretion ^61^, ^62^ was also reported to be transferred from the PM to ER in a signaling context at the level of MCS via Nir2 ^63^. Altogether, these findings highlight that ER-PM, or even ER-SG contacts as characterized here, could also act as a platform for lipid to fuel the requirements of the exocytic machinery. In line with this model, our recent data using novel clickable PA synthetic analogues identified lipid transfer proteins, typically enriched at MCS, as interacting partners of PA when chromaffin cells were stimulated for exocytosis ^64^. Our present data extends this concept by suggesting that lipid transfer may occur at ER-PM-SG-contacts. Although not directly addressed here, such lipid exchanges might influence the composition and curvature of the granule and plasma membranes and/or recruit specific cytoplasmic effectors (which might also be other tethers) further modulating their fusion competence. This lipid transfer could also be relevant following fusion to reshape typical lipid composition of SG and the PM after exocytosis, thus maintaining homeostasis of different membrane compartments involved in neurosecretion.

Another important implication of our work is the discovery of tripartite ER–PM–SG assemblies. These structures bring together three critical players of the exocytic machinery within nanometric distances, thus forming barriers for calcium diffusion across the whole cytoplasm. This may form privileged microdomains where calcium influx through STIM1/Orai1 channels and lipid transfer to the plasma membrane create a permissive environment for granule fusion. Such an organization could be particularly advantageous during sustained or repetitive stimulation, when secretion must be maintained despite the risk of local depletion of calcium or lipids. Interestingly, similar principles may apply to synaptic transmission, in which neurotransmitter release depends mainly on rapid Ca²⁺ influx through voltage-gated Ca²⁺ channels. Recent evidence indicates that axonal ER modulates synaptic transmission by regulating presynaptic Ca²⁺ dynamics ^65^. ER Ca²⁺ release or depletion enhances spontaneous and evoked neurotransmitter release, likely via ER-PM contact sites. STIM proteins may detect ER Ca²⁺ decreases and adjust neurotransmitter release by modulating voltage-gated Ca²⁺ channels at the PM ^66^. However, how synaptic ER Ca²⁺ uptake during neuronal activity modulates transmission remains unclear and still requires further investigation.

The identification of ER–granule and tripartite contact sites has broad implications for secretory physiology. In the adrenal medulla, modulation of ER–granule coupling may directly regulate catecholamine release during the stress response. Similar ER–vesicle contacts may operate in neurons, where their dysregulation could contribute to neurological disease^67^. The presence of Orai1 on secretory granules further suggests relevance for immune and endocrine systems. Orai1-mediated store-operated calcium entry is essential for cytokine secretion in T cells ^68^ and contributes to insulin release in pancreatic β-cells ^69^. Whether Orai1 similarly localizes to secretory vesicles in these contexts remains to be determined, but such localization would support ER–vesicle calcium coupling as a general mechanism, with disease-associated STIM1 or Orai1 mutations potentially impairing vesicular calcium homeostasis and exocytosis across tissues ^70^.

In summary, our findings uncover a novel role of the cortical ER as an active and versatile organizer of the exocytic machinery in neuroendocrine cells. By engaging in direct contacts with SG and forming tripartite assemblies with the PM, the ER emerges as a central integrator of calcium dynamics, lipid transfer, and vesicle maturation. These MCS could be especially relevant, during periods of high demand, when secretion is maintained over longer periods. Indeed, these contacts extend cellular control on local calcium buffering and lipid turnover to maintain secretion. Future work will be needed to dissect how these MCS are dynamically regulated by changes in tethering proteins, lipid composition, and post-translational modifications, and how such regulation shapes the efficiency and plasticity of secretory responses. Beyond chromaffin cells, these principles may apply broadly across secretory systems, with significant implications for endocrine regulation, neuronal communication, and disease ^67^.

## Materials & Methods

### Antibodies and reagents

The following primary antibodies were used in immunofluorescence experiments: anti-calnexin (rabbit, AB22595, Abcam, 1:200 dilution for fluorescent labelling and 1:100 for immunogold gold labelling); anti-Orai1 (rabbit, 13130-1-AP, Proteintech, 1:50 dilution), anti-STIM1 (mouse, sc-166840, Santa Cruz biotechnology, 1:100 dilution), anti-SNAP 25 (mouse, clone SMI81, cat 836304, Biolegend, 1:200 dilution), anti-DBH (mouse, clone 4F10.2, MAB 308, Millipore, 1:200 dilution, rabbit, ID: 22806, Immunostar, 1:500 dilution), anti-E-Syt (rabbit, HPA016858, Sigma Aldrich, 1:100 dilution), anti-PIP2(Mouse, ab11039, Abcam, 1:50 dilution) and Biotin XX phalloidin (B7474, Thermo Fisher Scientific Inc., 1:100 dilution).

The following fluorescent secondary antibodies were used in immunofluorescence microscopy: goat anti- -mouse and -rabbit IgG (H+L) *highly cross-adsorbed* conjugated to Alexa Fluor® -488, -555, or -647 (ThermoFisher, all used at a 1:1000 dilution). Hoechst 33342 was used to label DNA. Secondary antibodies and streptavidin coupled gold particles (1:100 dilution) were from Aurion (Wageningen, NL) and Thermo Fisher Scientific Inc. (Waltham, MA, USA) respectively. Nicotine (N3876MG), BTP2 (203890-M) and solid salts were obtained from Sigma Aldrich. Hoechst 33258 (461061) was from Dutscher.

### Chromaffin cells culture

Bovine adrenal glands were sampled from the Haguenau municipal slaughterhouse in the framework of authorization n°67-482-1683 granted by the lower-Rhin departmental population protection direction for sampling and usage of animal by-products. They were placed in fresh calcium-free Krebs solution (NaCl 154 mM, KCl 5.6 mM, NaHCO_3_ 3.6 mM, 5.6 mM glucose and 5 mM HEPES, pH 7.4) until the start of the culture. Chromaffin cells were isolated from fresh bovine adrenal glands by retrograde perfusion with collagenase, purified on self-generating Percoll gradients and maintained in culture as previously described ^64^. To induce exocytosis, on the day of experiments, chromaffin cells were washed twice with Locke’s solution (140 mM NaCl, 4.7 mM KCl, 2.5 mM CaCl_2_, 1.2 mM KH_2_PO_4_, 1.2 mM MgSO_4_, 11 mM glucose, 0.56 mM ascorbic acid, 0.01 mM EDTA and 15 mM HEPES, pH 7.5) then stimulated with Locke’s solution containing 20 µM nicotine or 59 mM K^+^ depolarizing solution (85.7 mM NaCl, 59 mM KCl, 2.5 mM CaCl_2_, 1.2 mM KH_2_PO_4_, 1.2 mM MgSO_4_, 11 mM glucose, 560 µM ascorbic acid, 10 µM EDTA and 15 mM HEPES, pH 7.2) as specified in figure captions.

### DNA constructs and cell transfection

Plasmids WT STIM1-Ac GFP and L251S STIM1-Ac GFP were described by Korzeniowski ^35^. Genetic modification of chromaffin cells was performed under the GMO authorization DUO 117 granted by the French ministry of research and higher education to the INCI UPR3212. Plasmids (5 µg) were transfected into chromaffin cells (5 x 10^6^ cells) by nucleofection, immediately following culture, using the primary basic nucleofector kit for primary neurons (Amaxa Nucleofactor systems, Lonza, Levallois, France) according to the manufacturer’s instructions. Electroporated cells were immediately recovered in warm antibiotics and antimitotic free culture medium and plated onto fibronectin-coated glass coverslips. Both antibiotics and antimitotic were added back to the cells 5 h after the transfection procedure and medium was changed the following day. Experiments were performed 3 to 4 days after transfection.

### Immunofluorescence and confocal microscopy

For immunocytochemistry, chromaffin cells were grown on fibronectin-coated glass coverslips, fixed and labelled as described previously ^19^. The transient accessibility of DBH to the PM of chromaffin cells was tested by incubating cells for 3 min in stimulation solution (15 µM nicotine of 59 mM K^+^) containing anti-DBH antibodies diluted to 1:100.

Labelled cells were visualized using a Leica SP5II confocal microscope equipped with a planapo oil immersion objective (63X, n.a. = 1.4, pinhole set to 0.68 Airy unit). The proportion of colocalization was estimated from double-labelled pixels using ICY software (http://icy.bioimageanalysis.org).

### Chromaffin cells slice and PM sheet preparation and scanning for electron microscopy

Resting and stimulated chromaffin cells were fixed with 2% glutaraldehyde in 0.1 M phosphate buffer for 30 min at room temperature. After fixation, the samples were washed in phosphate buffer for 2 h. Cells were postfixed (0.8% potassium ferrocyanide to enhance membrane contrast ^71^ and 0.5% OsO_4_ in ultrapure water). Finally, samples were dehydrated in graded ethanol series and embedded in Embed 812-Araldite mix (EMS). Ultrathin sections were cut (60 to 80 nm) with an ultramicrotome (Leica), stained with lead citrate (Delta Microscopies), and examined by TEM (Hitachi H7500 equipped with an AMT Hamamatsu digital camera). Cytoplasmic face-up membrane sheets were prepared and processed as previously described ^30^. Briefly, carbon-coated Formvar films on nickel electron grids were inverted onto unstimulated or stimulated chromaffin cells. To prepare membrane sheets, a pressure was applied to the grids for 20 s, then grids were lifted so that the fragments of the upper cell surface adhered to it, which required less than 30 s. Immediately after cell stimulation and sheet preparation, these membrane portions were fixed in 2% paraformaldehyde in PBS for 10 min at 4°C. After blocking in PBS with 1% BSA and 1% acetylated BSA, immunolabelling was performed and revealed with 10 nm, 15 nm and 25 nm gold particles-conjugated secondary antibodies. These membrane portions were fixed in 2% glutaraldehyde in phosphate buffer 0.1M, postfixed with 1% OsO_4_, dehydrated in a graded ethanol series, treated with hexamethyldisilazane (Sigma-Aldrich, St. Louis, MO, USA), air-dried and observed using a Hitachi 7500 TEM ^26^. Granules were counted on random TEM images of PM sheets (n= 25 to 50 images of 45 µm^2^ of PM). The data are expressed as number of granules per µm^2^.

### Electron tomography and image processin**g**

Tilt series were acquired on a JEOL JEM-2100 transmission electron microscope operated at 200 kV. Data were automatically acquired using Digital Micrograph software. Typically, the tilt ranged between - 60 and + 60 with 2° angular increments. Images were recorded at a nominal magnification of 10,000 x on a Gatan Ultrascan 2K×2K CCD camera with defocus set to − 2 μm. ^26^. Alignments of raw tilt series, using anti-calnexin immunogold particles as fiducial markers, were computed with IMOD software package ^72^.Tomographic reconstruction was carried out using either weighted back projection for cytoplasmic face-up membrane sheets or SIRT (Simultaneous Iterative Reconstruction Technique) for ultrathin sections. The PM, secretory granules and ER were manually segmented using drawing tools in IMOD.

### Proximity ligation assay (PLA)

In situ PLA kit Duolink (DUO92101-1KT, Sigma-Aldrich) was used to detect the interaction of Orai1 and STIM1 according to the manufacturer’s instructions. Briefly, bovine chromaffin cells were fixed and permeabilized. Next, cells were blocked with blocking solution and incubated with rabbit anti-Orai1 and mouse anti-STIM1 (1:50, each) antibodies overnight at 4 °C. After 2 washes, cells were incubated with Duolink secondary antibodies conjugated with oligonucleotides (anti-rabbit PLA probe Plus and anti-mouse PLA probe Minus) for 1 h at 37 °C. Then, cells were incubated with a ligation solution containing oligonucleotides and the ligase. The amplification products were detected with red fluorescence-labeled complementary oligonucleotide detection probes. Samples were mounted with a medium containing 4′,6-diamidino-2-phenylindole (DAPI) nuclear stain. The fluorescence signals (positive signals: red fluorescent puncta) were visualized and imaged using a TCS SP5II confocal microscope (Leica, Germany) ^73^. After acquisition of a Z-stack (pinhole set to 1 Airy unit, step size of 0.3 µm), puncta present on the maximum-intensity Z-projection were counted manually.

### Carbon fiber amperometry

Bovine chromaffin cells were washed with Locke’s solution and processed for catecholamine release measurements by amperometry as previously described ^74^. Cells were used 3 to 4 days after culture washed twice with ascorbic acid free Locke’s solution (140 mM NaCl, 4.7 mM KCl, 2.5 mM CaCl_2_, 1.2 mM KH_2_PO_4_, 1.2 mM MgSO_4_, 11 mM glucose, 0.01 mM EDTA and 15 mM HEPES, pH 7.5) and recordings were performed in the same medium over the next 60 min. For experiments with BTP2 or DMSO (control), the molecule was added directly in the incubation medium 10 min before starting recordings and left in the medium during recordings. A carbon fiber electrode of 5 μm diameter (ALA Scientific Instruments) was held at a potential of +650 mV compared to the reference electrode (Ag/AgCl) and approached closely near the recorded cell. Secretion of catecholamines was induced by a 10 s pressure ejection of a 100 mM ascorbic acid free K^+^ solution (44 mM NaCl, 100 mM KCl, 2.5 mM CaCl_2_, 1.2 mM KH_2_PO_4_, 1.2 mM MgSO_4_, 11 mM glucose, 10 µM EDTA and 15 mM HEPES, pH 7.2) with a micropipette (Femtotips®, Eppendorf) positioned approximately 10 μm from the cell and recorded over 60 s. For transfected cells, GFP positive cell were identified thanks to a pE-4000 illumination system (CoolLED) with a 490 nm LED emission. PolyScan2 and Micro-Manager 1.4 were used to control the LED. The amperometric recordings were performed with an AMU130 amplifier (Radiometer Analytical), calibrated at 5 kHz, and digitally low-pass filtered at 1 kHz. Analysis was performed as previously described with a macro (laboratory of Dr. R. Borges; http://webpages.ull.es/users/rborges/) written for Igor software (WaveMetrics), allowing automatic spike detection and extraction of spike parameters ^75^. Only cells having produced at least 5 spikes were included in the analysis, except if the inhibition was too strong (BTP2, and L251S STIM mutant groups), in which case the proportion of responding cells in the control group was applied to treated or mutant groups to include or exclude cells.

The individual spike parameters analysis was restricted to spikes with amplitudes higher than 5 pA, which were considered as exocytic events ^76, 77^. All spikes detected by the program were visually inspected. Overlapping and aberrantly shaped spikes were counted in the total number of events but discarded for individual spike parameters analysis. Quantal size (spike charge, Q) of each individual spike was measured by calculating the spike area above the baseline, the spike area being defined as the time integral of each transient current. Imax was defined as the height of each spike, half-width as the width of each spike at half its height (T_1/2_) and T_peak_ as the spike rise time (Figure 5C).

### Statistical analysis

Statistical analysis and graphical representations were performed on the Graphpad Prism software. As specified in figure legends, group data are presented as mean (± SEM) and each point represent a single measurement. Statistical differences were established using a Mann-Whitney test, or a Kruskal-Wallis test followed by Dunn’s multiple comparisons post-hoc test. Asterisks on plots indicate statistical significance (*: p < 0.05; **: p < 0.01; ***: p < 0.001 and ****: p < 0.0001).

## Abbreviations

BTP2: 3,5-BisTrifluoromethyl Pyrazole
Ca^2+^: Calcium Ion
DBH: Dopamine Beta-Hydroxylase
ER: Endoplasmic Reticulum
E-Syts: Extended Synaptotagmins
MCS: Membrane Contact Sites
Orai1: Calcium Release-Activated Calcium Modulator 1
PA: Phosphatidic Acid
PIP_2_: PhosphatidylInositol-4,5 –bisPhosphate
PLA: Proximity Ligation Assay
PM: Plasma Membrane
PS: Phosphatidyl Serine
SERCA: SarcoEndoplasmic Reticulum Ca^2+^-ATPase
SG: Secretory Granule
SNAP25: SyNaptosomal Associated Protein of 25 kDa
STIM1: STromal Interaction Molecule 1
TEM: Transmission Electron Microcopy
VAMP: Vesicular Associated Membrane Protein
VAP: VAMP-Associated Proteins

## Acknowledgments

We thank Marek Korzeniowski and Tamas Balla (NICHD, NIH, USA) for sharing the STIM-1 plasmids and Stephane Ory for his help in using the ICY software. The In Vitro Imaging Platform-Strasbourg (CNRS UAR3156) is member of the national infrastructure France BioImaging supported by the French National Research Agency (ANR-10-INSBS-04). Part of this work was financially supported by the Agence Nationale pour la Recherche ANR-19-CE16-0012 to S.G., ANR-19-CE44-0019 and ANR-24-CE44-1240 to NV . INSERM is providing salary to S.C.G., S.G., and N.V. We acknowledge the municipal slaughterhouse of Haguenau (France).

## Author contributions

F.D. conducted electron tomography and detailed analysis. L.S. performed carbon fiber amperometry experiments and analysis. A.W. conducted bovine chromaffin cell cultures. C.R. A-M H. performed membrane sheets, cell section labellings and associated TEM imaging. S;H. performed Ca^2+^ imaging. S.C-G formulated the original hypothesis, performed immunofluorescent labellings, PLA, and associated confocal imaging. A.W., N.V., S.G. and S.C-G. drafted the manuscript. N.V., S.G. and S.C-G supervised the study. All authors discussed the results and contributed to the manuscript revision.

## Data availability

The data that support the findings of this study are available from corresponding authors upon reasonable request.

## Additional information

Supplemental Figures 1, 2

Tables I, II

Supplemental Video 1 - Tomogram of an unstimulated chromaffin cell section.

Supplemental Video 2 - 3D surface rendering of the segmented subtomogram from an unstimulated chromaffin cell.

Supplemental Video 3 - Tomogram of a nicotine-stimulated chromaffin cell section.

Supplemental Video 4 - 3D surface rendering of the segmented subtomogram from a stimulated chromaffin cell.

Supplemental Video 5 - Tomogram of calnexin-positive structures associated with secretory granules (SGs) docked at the plasma membrane (PM), corresponding to Fig. 4a.

Supplemental Video 6 - Tomogram of calnexin-positive structures associated with SGs docked at the PM, corresponding to Fig. 4b.

Supplemental Video 7 - Tomogram of calnexin-positive structures associated with SGs docked at the PM, corresponding to Fig. 4g.

Supplemental Video 8 - Tomogram of calnexin-positive structures associated with SGs docked at the PM, corresponding to Fig. 4h.

Supplemental Video 9 - 3D animation of Z-projected in situ PLA signals showing STIM1–Orai1 interaction in unstimulated chromaffin cells.

Supplemental Video 10 - 3D animation of Z-projected in situ PLA signals showing STIM1–Orai1 interaction in stimulated chromaffin cell.

