## Supplemental figures for "Endoplasmic reticulum membrane contact sites coordinate exocytic site assembly and activity in neuroendocrine cells"

### Supplemental Fig. 1

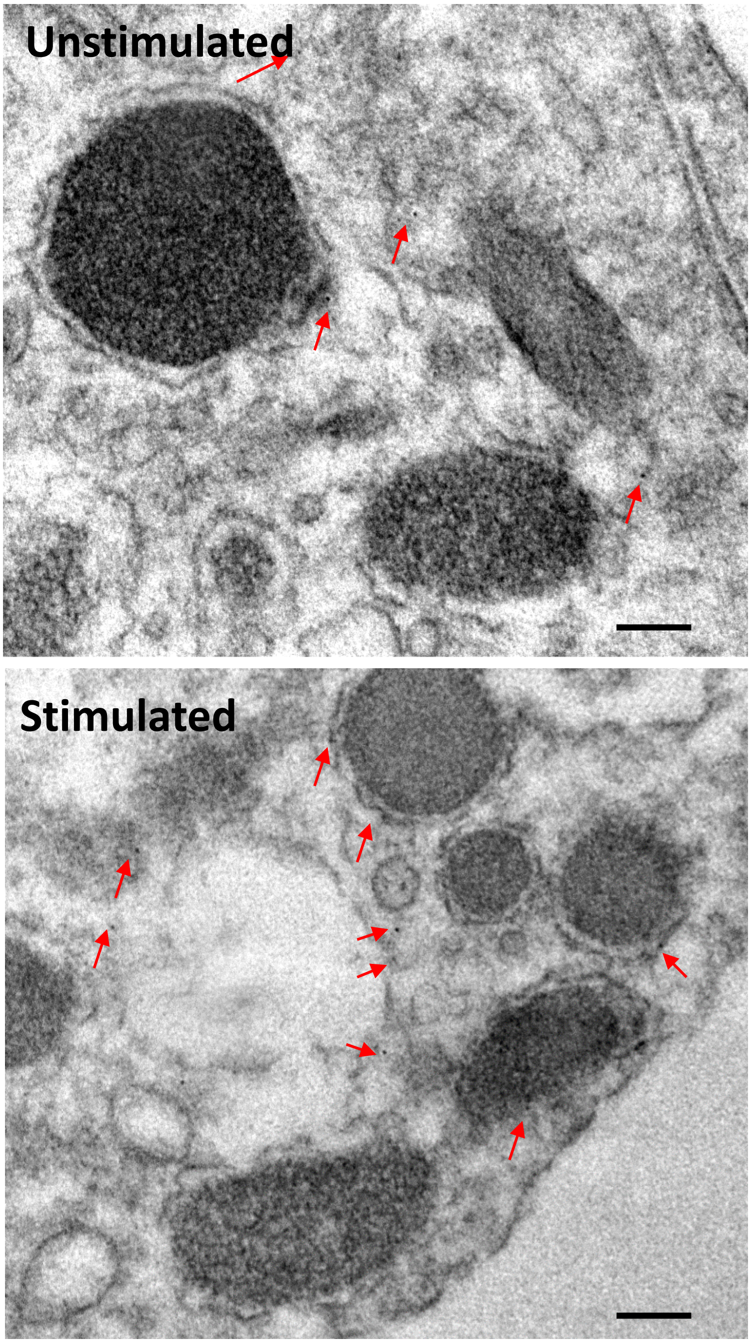


**Figure S1: Labeling of calnexin in unstimulated and stimulated chromaffin cells ultrathin sections.** Representative transmission electron microscopy images of chromaffin cells subjected to pre-embedding immunogold labeling. Cells were fixed with 4% paraformaldehyde and 0.1% glutaraldehyde, incubated with sodium borohydride, and blocked with BSA. Calnexin was detected using a rabbit anti-calnexin primary antibody followed by 6-nm gold-conjugated $F(ab')_2$ fragments of goat anti-rabbit IgG. Samples were post-fixed with glutaraldehyde and osmium tetroxide, dehydrated, embedded in Embed 812, and sectioned for transmission electron microscopy. Control experiments included omission of the primary antibody, which resulted in negligible nonspecific labeling. Gold particles (arrowheads) indicate the localization of calnexin at the ultrastructural level. Scale bars, 100 nm.

### Supplemental Fig. 2

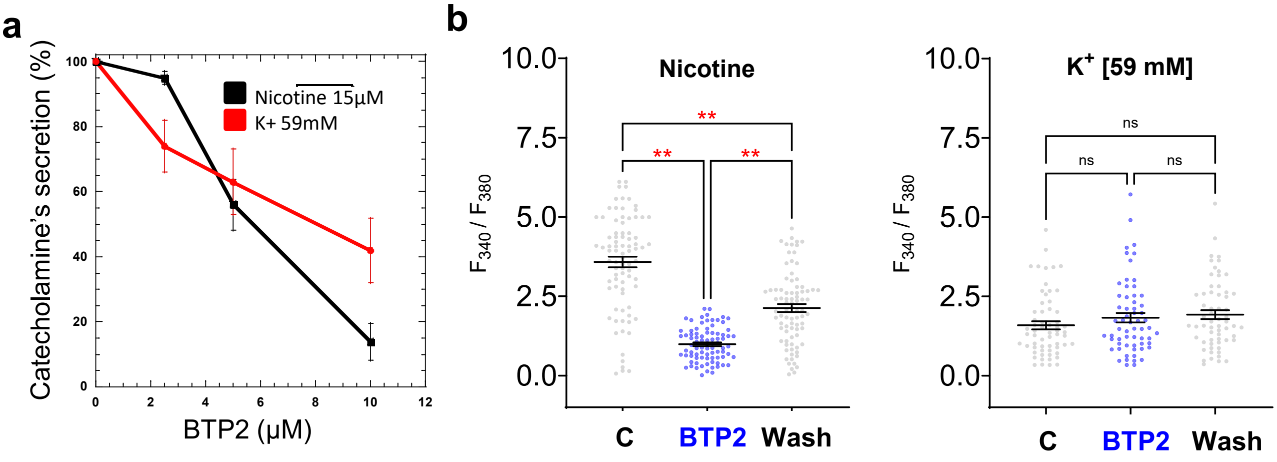


**Figure S2 : BTP2 inhibits catecholamine release and nicotine-evoked $Ca^{2+}$ signals in chromaffin cells.**

**(A**) Chromaffin cells were washed three times with Locke's solution and incubated for 30 min with BTP2 at the indicated concentrations. Cells were then kept at rest or stimulated for 10 min with 59 mM $K^+$ or 15 µM nicotine in the continued presence of BTP2. Catecholamine release was quantified fluorimetrically as described (Tahouly et al., 2021, Methods Mol Biol. 2233:169-179). Briefly, 20 µl of supernatant were oxidized with sodium acetate (1 M, pH 6) and $K_3Fe(CN)_6$, then converted to adrenolutin using NaOH/ascorbic acid, and fluorescence was measured at λ_ex 430 nm / λ_em 520 nm (LB940 Mithras). Values are expressed as % net secretion. Data show a representative experiment from six independent assays (mean ± SEM).

**(B)** Fura-2 $Ca^{2+}$ imaging was performed 3 d after seeding. Cells were stimulated three times with nicotine (15 µM) or $K^+$ (59 mM; 5 s pulses, 600 s interval), with BTP2 (10 µM) co-applied during the second stimulation. Peak F340/F380 ratios were quantified as described (Eersapah et al., 2019, Purinergic signaling, 15(3): 403-420). A repeated-measures ANOVA revealed a strong inhibitory effect of BTP2 on nicotine-evoked $Ca^{2+}$ signals ($F(2,164) = 262.3$, $p < 0.0001$, 83 cells, 6 cultures), with partial recovery after washout ($p < 0.0001$ vs initial response). By contrast, $K^+$-evoked $Ca^{2+}$ responses were unaffected ($F(2,120) = 2.33$, $p = 0.10$; 61 cells, 4 cultures).
